# Relating neural oscillations to laminar fMRI connectivity

**DOI:** 10.1101/2020.09.18.303263

**Authors:** René Scheeringa, Mathilde Bonnefond, Tim van Mourik, Ole Jensen, David G. Norris, Peter J. Koopmans

## Abstract

Laminar fMRI holds the potential to study connectivity at the laminar level in humans. Here we analyze simultaneously recorded EEG and high resolution fMRI data to investigate how EEG power modulations, induced by a task with an attentional component, relate to changes in fMRI laminar connectivity between and within brain regions. Our results indicate that our task induced decrease in beta power relates to an increase in deep-to-deep layer coupling between regions and to an increase in deep/middle-to-superficial layer connectivity within brain regions. The attention-related alpha power decrease predominantly relates to reduced connectivity between deep and superficial layers within brain regions, since, unlike beta power, alpha power was found to be positively correlated to connectivity. We observed no strong relation between laminar connectivity and gamma band oscillations. These results indicate that especially beta band, and to a lesser extent alpha band oscillations relate to laminar specific fMRI connectivity. These differential effects for the alpha and beta bands suggest a complex picture of possibly co-occurring neural processes that can differentially affect laminar connectivity.

## INTRODUCTION

Investigating directional laminar connectivity during task conditions has so far remained primarily the domain of invasive electrophysiological investigations in animals (van Kerkoerle et al 2014). Over the past decade, the development of high-resolution fMRI has made it possible to measure laminar level activity in humans non-invasively (Finn et al 2019, Huber et al 2017, Kok et al 2016, Koopmans et al 2010, Lawrence et al 2018, Muckli et al 2015, Polimeni et al 2010, Sharoh et al 2019, Siero et al 2011). Laminar specific anatomical connections directly relate to feedforward and feedback projections between regions at different levels in the cortical hierarchy (Douglas & Martin 2004, Markov & Kennedy 2013). These projections have been related to frequency-specific directional connectivity derived from electrophysiological measures. Recently, Sharoh et al. (2019) demonstrated that laminar specific feedforward and feedback connections of the visual word form area with other regions can be studied with cortical depth resolved fMRI. Since frequency-specific electrophysiological synchronization has been directly related to laminar specific feedforward and feedback projections, this raises the question of how frequency-specific neural synchronization relates to laminar level fMRI connectivity (Scheeringa & Fries 2017). Several frameworks for cortical processing explicitly link frequency-specific neural synchronization to laminar specific feedforward and feedback projections between brain regions (Bastos et al 2012, Bonnefond et al 2017, Fries 2015). Laminar level fMRI connectivity could provide a unique opportunity to study such feedforward and feedback projections non-invasively in humans that can contribute to testing theoretical frameworks like these. For this, it is essential to first establish how neural oscillations in separate frequency bands relate to laminar connectivity. In an exploratory re-analysis of simultaneously recorded laminar fMRI-EEG data during an attention-demanding task (Scheeringa et al 2016) we therefore investigate how modulations of neural oscillations relate to changes in connectivity between regions in the early visual cortex.

Invasive recordings in animals have linked feedforward connectivity to inter-regional synchronization in the gamma and theta bands, while feedback has been linked to alpha and beta band synchronization (Bastos et al 2012, Bastos et al 2015, Bosman et al 2012, van Kerkoerle et al 2014). Anatomically, these gamma/theta band feedforward connections are thought to originate from superficial (layers II/III) in lower order regions and target mainly middle layers (layer IV) in higher order regions. Feedback related alpha and beta band activity is thought to be conveyed from deep layers in higher order regions (layers V/VI) (Buffalo et al 2011, Fries 2015, van Kerkoerle et al 2014) to layers outside layer IV in lower order regions, although for nearby regions feedback connections between superficial layers (layers II/III) have also been identified (Markov et al 2013). Although true anatomical layers can at the moment not be uniquely identified and measured with high resolution fMRI, the sub-millimeter resolution that can now be achieved does allow for depth resolved measures of the hemodynamic response (roughly 3-4 independent observations over the width of the cortex) at a similar spatial scale to the underlying three cortical ‘super’ layers defined as infra-granular, supra-granular and granular. Laminar fMRI therefore allows us to study connectivity between various cortical depths non-invasively (Huber et al 2020, Huber et al 2017, Sharoh et al 2019, Wu et al 2018) and when combined with electrophysiological measures like EEG we can relate these effects to frequency specific neural synchronization. Here we implemented this combination by first computing individual task effects in cortical depth dependent correlation-based connectivity. Subsequently we correlated this task effect over subjects with task effects in frequency specific EEG power bands.

In the original analyses of the data used here (Scheeringa et al 2016), we demonstrated that the alpha, beta and gamma band effects observed in this task correlated to the BOLD effect in different cortical layers. Alpha oscillations were found to correlate negatively with both deep and superficial layers, while beta oscillations were found to correlate negatively only with deep layer BOLD. For the gamma band a positive correlation was found in middle and superficial layers. These findings were largely in line with the likely laminar origins and function of these rhythms, suggesting a link between laminar BOLD and neural synchrony related feedforward and feedback projections between brain regions (Bastos et al 2015, van Kerkoerle et al 2014). Crucially, the experimental paradigm task included a crude attention manipulation (attention ON vs. Attention OFF; see Fig. 1A) that forms the basis for the present contribution. Recent publications on laminar fMRI have demonstrated the possibility of performing laminar level fMRI connectivity analyses (Huber et al 2020, Sharoh et al 2019) for both intra- and inter-regional connectivity. This led us to revisit our previously recorded data in order to explore whether not only laminar specific BOLD amplitude, but also laminar fMRI connectivity relates to frequency specific EEG power. Specifically, by correlating task effects in laminar connectivity within and between regions in the visual cortex (V1, V2, V3), with task effects in alpha, beta and gamma bands (see Fig. 1B-C for the original findings for this task contrast).

**Fig. 1:**
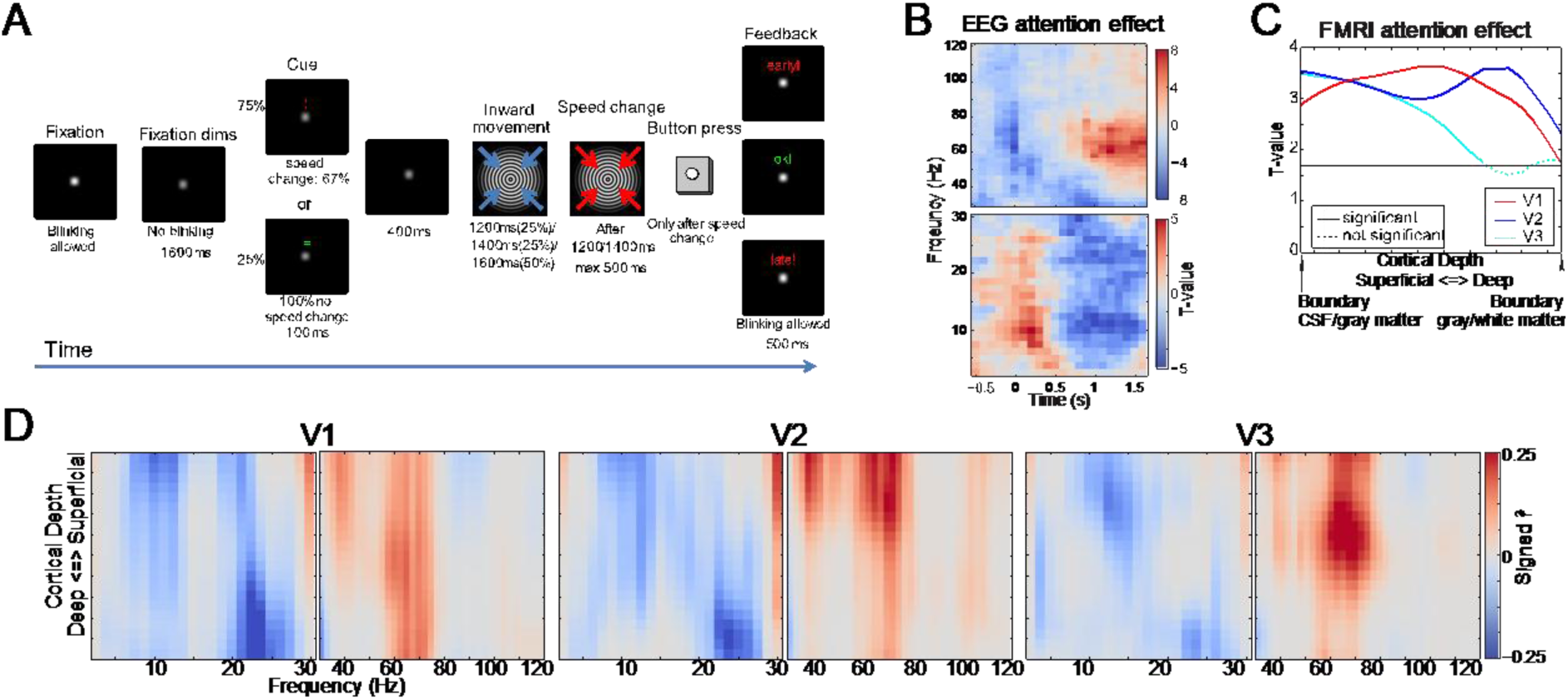
Experimental paradigm and attention effects. A) Schematic representation of a trial of the experimental paradigm. Subjects were instructed to focus on a fixation dot, that would dim 1000 ms before trial onset and indicated that subjects were not to blink anymore. At the trial onset subjects would see a cue for 100 ms indicating whether a speed increase in the inward contractions of the grating would likely occur (75% of the trials) or certainly not occur (25% of the trials). If so, these changes would start 500 ms after the onset of the cue. If a speed change was cued, it would occur in 67% of the trials, and not occur in 33 % of the trails. The trials where a speed-increase was cued but did not occur (attention ON condition) and the trials where the cue indicated a speed change would definitely not occur (attention OFF condition) are the basis of the work presented here (attention contrast). B) Time-frequency representation of the t-values for the attention contrast in EEG power which was computed as the log-transformed ratio of the power in both conditions. Time is relative to the onset of visual stimulation. C) T-values across the cortical depth of the cortex for the attention contrast for the cortical depth resolved BOLD signal for V1, V2 and V3. The BOLD signal was estimated for 21 points across the width of the cortex, where the boundary between CSF and Gray matter coincides with the left limit of the plot and the border between gray and white matter with the right limit. D) Signed square correlation between the attention effect in the EEG power averaged over 0.6-1.6 s during the visual stimulation and the attention effect in depth resolved BOLD for V1, V2, V3. Here, the boundary between CSF and Gray matter coincides with the top of the y-axis and the border between gray and white matter with the bottom of the y-axis. Frequency is plotted on the x-axis, separately for low (<30 Hz) and high (40-120 Hz) frequencies due to the difference in the ranges of the frequency-smoothing for the two frequency ranges (<30 Hz): +/-2.5Hz; 30-120 Hz: +/-10 Hz). The results depicted in panels B-D were previously reported in Scheeringa et al. (2016) and all panels were adapted from this previous work.

The results from the new analyses presented here should be interpreted as an exploration of how laminar fMRI based estimates of inter- and intra-regional coupling relate to neural oscillations. We chose a paradigm that was known from previous studies using MEG and EEG (Hoogenboom et al 2010, Hoogenboom et al 2006, Muthukumaraswamy & Singh 2013) to induce changes in multiple frequency bands in the early visual cortex. Furthermore, the EEG responses are highly similar to those invasively recorded in animals from visual regions in tasks investigating selective attention (Fries et al 2008). Together this gives us confidence that the signals we measure originate from early visual cortex, although the source location cannot be accurately be estimated from our data due to the relatively low number of channels and the larger spatial imprecision of EEG compared to MEG and intracranial recordings. The crude task modulation added to the paradigm (attention On versus Off) was introduced in the first place to induce meaningful variation over subjects in task effects across the frequency bands modulated by the visual stimulation, as well as across all layers of the cortex. It was not intended to investigate specific individual processes such as prediction, attention or arousal. In this study we make use of the variation in task effect over subjects, by correlating the task effects in alpha, beta and gamma band power to the task effects in measures of inter- and intra-regional coupling over subjects.

The frequencies investigated have been previously directly linked to specific processes (Bastos et al 2012, Fries 2015, Jensen & Mazaheri 2010, Klimesch et al 2007). Within paradigms addressing selective attention and prediction, gamma band oscillations are predominantly observed in the superficial layers and have been associated with attention modulated gain of the feed forward stream (Bastos et al 2015, Bosman et al 2012, Fries 2015, Michalareas et al 2016, Roberts et al 2013) and, within a predictive coding framework the feed forward projection of prediction errors (Bastos et al 2020, Bastos et al 2012). Within this framework alpha/beta oscillations are thought to reflect feedback projections that carry predictions from higher order to lower order regions (Bastos et al 2020, Bastos et al 2012). Others have associated alpha and beta oscillations with inhibition of task irrelevant regions whose activity could otherwise interfere with task performance (Jensen & Mazaheri 2010, Klimesch et al 2007, Zumer et al 2014). All these perspectives explicitly or implicitly rely on (changes in) laminar specific neural coupling within and between brain regions. Laminar level fMRI connectivity holds the unique key to measure such coupling non-invasively in humans (Scheeringa & Fries 2019). The present work links laminar fMRI connectivity to such neural oscillation-based models of cognitive brain function in two ways. First, we shed light on whether laminar fMRI-based connectivity (either alone or combined with EEG) can be used to address processes traditionally investigated with invasive electrophysiology. Second, the presence (or absence) of a relation between laminar connectivity (measured by fMRI) and neural oscillations (measured by EEG) is relevant for cognitive models that predict specific roles for neural oscillations in relation to laminar level coupling within and between brain regions.

## RESULTS

### Previous results

A full account of our initial analyses and results can be found in Scheeringa et al. (Scheeringa et al 2016). In panel A of Fig. 1 we depict the structure of the experimental paradigm, an inward contracting circular sinusoidal grating, where subjects have to react to a speed increase in this contraction with a button press. Crucial for the analyses presented here, a cue indicated this was likely to occur (75% of the trials, 67% cue validity) or would not occur at all (25% of the trials, 100% cue-validity). The analysis presented here pertains to the two conditions where no button-press was required; the “attention ON” condition where a speed-change was cued but did not occur, and the “attention OFF” condition where no speed change was cued. This paradigm and task manipulation were chosen because they induce decreases in alpha and beta power of the EEG and increases in gamma band power relative to baseline, as well as in the attention ON condition compared to the attention OFF condition. This allows us to investigate the relationship of oscillatory power in different frequency ranges with laminar level BOLD activity (Scheeringa et al 2016) and connectivity (presented here) in the same task. As a consequence, our task modulation was not a clean modulation of attention and should not be interpreted as such, since relevant cognitive processes like arousal and the predictability of the speed change differed between the conditions.

In Panels 1B-D we depict the relevant results from this attention contrast taken from our previous work. In panel B we show a time-frequency representation of the effects on EEG power. It shows that alpha and beta power in the last second of the trials are lower during “attention ON” trials than during “attention OFF” trials, while gamma power is higher during the “attention ON” condition. In Panel C we show stronger BOLD activation during this condition across all cortical depths in V1, V2 and V3. For the fMRI signal we estimated the BOLD signal for 21 points between pairs of vertices at the CSF/gray matter and gray/white matter boundaries. The analyses presented here were carried out on BOLD data averaged from the top 10% activated pairs of vertices in each region (calculated over attention conditions and cortical depth). This selects those parts of the regions that were active in response to the stimulus which spanned only a part of the visual field (changing the 10% threshold value to 5% and 25% did not yield a qualitatively different picture). Averaging over task activated regions in V1/V2/V3 is necessary to increase the signal to noise level, which is very low for the high-resolution individual voxels. The results for Panel C show that attention modulation leads to an increased BOLD response across all cortical depths in all three regions. In panel D we show the (signed squared) partial correlation between the attention effects in EEG power and those in laminar BOLD, while controlling for the effect of pupil diameter. These analyses show that the EEG alpha attention effect is negatively correlated with the BOLD attention effect in superficial layers for all regions, while the beta effect correlates negatively with the deep layer BOLD attention effect. For gamma we observed a positive correlation in middle/superficial depths for V2 and V3, and across all cortical depths for V1.

### Laminar Connectivity

Before we discuss the details of the analysis below, we summarize its rationale: we wish to relate (changes in) connectivity to neural oscillations. Typical connectivity analyses tend to condense a series of measurements (e.g. time series data) into a single value, which subsequently is hard to relate to anything else (e.g. EEG signals) over time. We therefore calculate a correlation *over subjects* between subject-specific fMRI connectivity measures and their EEG power counterparts. In summary: For each individual subject’s fMRI data we calculate the connectivity between the signals of two layers, separately for both the “attention ON” and “attention OFF” conditions. These two layers can be in the same region (Fig. 4) or in different regions (Fig. 3). We calculate a difference score for attention ON and OFF which is then taken as the attention effect in connectivity for that particular combination of layers and regions, and, importantly, for that specific subject. To establish the link with the EEG frequency bands, we subsequently assess the correlation across subjects of this subject-specific fMRI attention contrast value with the subject-specific attention effects in the EEG power for each frequency band. So functional connectivity is assessed at the subject level (i.e. one value per subject) whereas EEG and fMRI are linked at the group level. The above is repeated for all possible combinations of layers and regions. Results are displayed in a correlation matrix style with cortical depth on both axes. Interregional results are shown in Fig. 3 and intraregional results in Fig. 4.

**Fig. 2:**
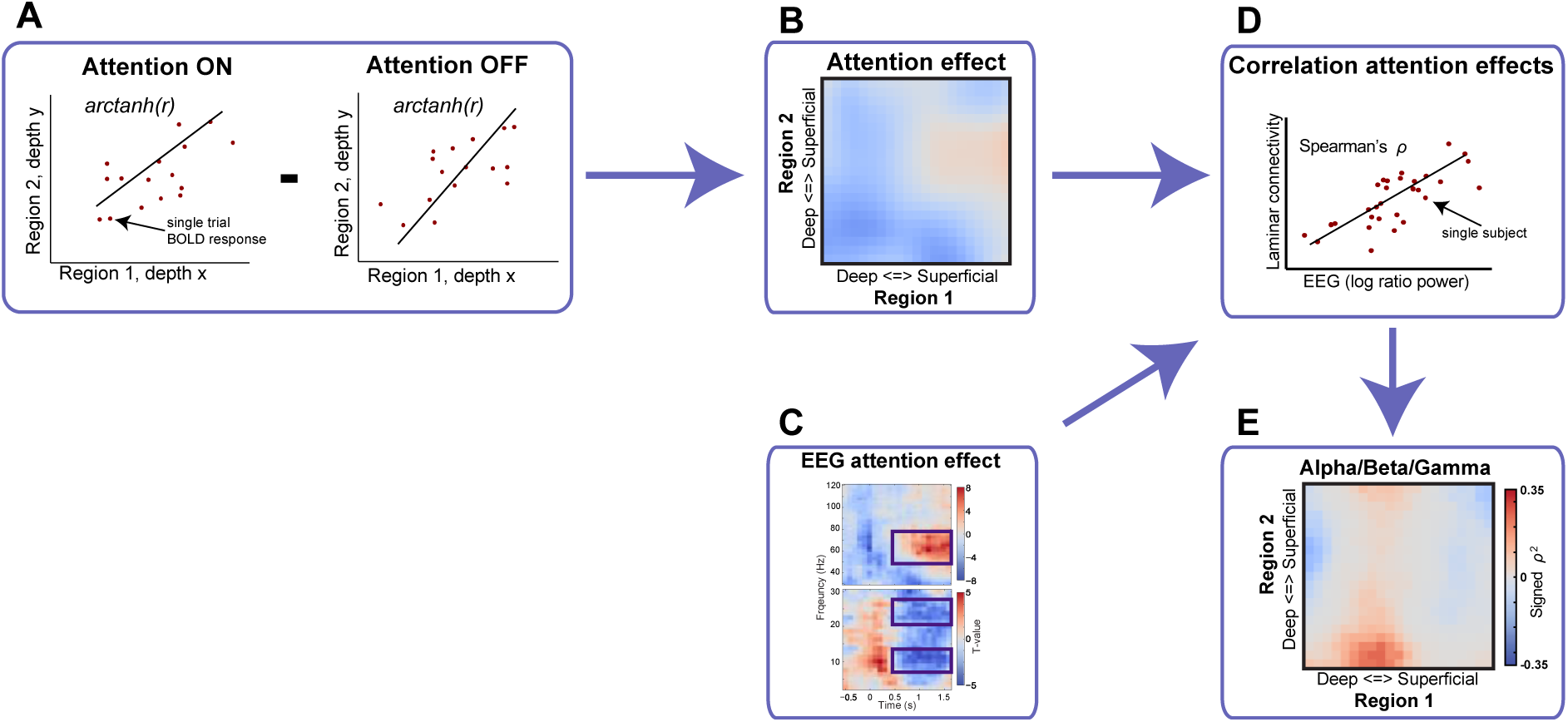
Schematic representation of the analysis pipeline. Laminar connectivity was computed as the Fisher’s z-transformation of the Pearson correlation (r) between the amplitude of single trial BOLD responses, which were obtained through deconvolution (panels A) for both conditions and separately for each block. The attention effect for a single subject, depicted for each depth combination between regions in panel B, is computed as the difference between these z-transformed correlations averaged over the three experimental blocks. The single subject attention effects in the alpha, beta and gamma bands (panel C) were correlated over subjects with the single subject attention effects in laminar connectivity (panel D) using Spearman’s rank-order correlation (ρ). After squaring and multiplication with the sign of the correlation, the laminar level relation of EEG frequency band with depth-resolved connectivity between region-pairs can be assessed separately for alpha, beta and gamma (panel E). For the analysis presented in the main article laminar connectivity was first averaged over the region pairs indicated in Fig. 3 & Fig. 4 before the attention effect was calculated. For the results in supporting Figs. S1-S4 this attention effect was calculated for individual region-pairs.

**Fig. 3:**
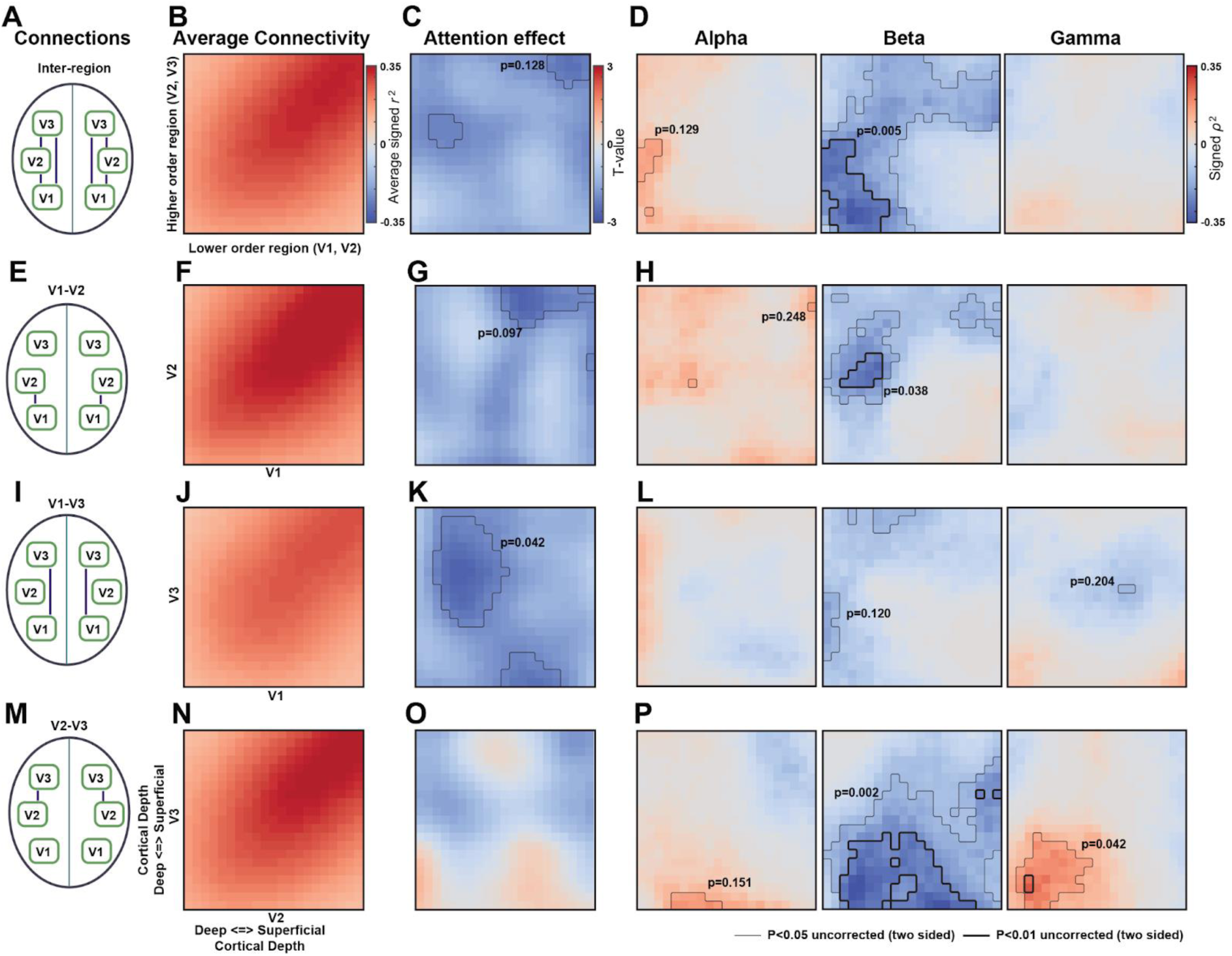
Relation between EEG power and inter-regional laminar fMRI connectivity. The top row (panels A, B, C & D) depicts the results for inter-regional connectivity (our main analysis). The three rows below depict the results for the individual connection pairs V1-V2 (panels E, F, G & H), V1-V3 (I, J, K & L) and V2-V3 (M, N, O & P). The left column (A, E, I & M) illustrates for each row the connections over which connectivity was averaged, and the results in the panels to the right apply. For these panels the lower order region in the visual hierarchy is plotted on the x-axis, and the higher order region on the y-axis. On the x-axis, deep to superficial cortical depths are depicted from left to right, while on the y-axis they are depicted from bottom to top. In the second column (B, F, J & N) depicts the average connectivity over attention ON and attention OFF conditions, calculated as the average over runs conditions and subjects of the squared Pearson correlation multiplied by the sign of the correlation. The third column (C, G, K & O) depicts the task effect on laminar connectivity expressed in t-values. The three right most columns (panels D, H, L & P) show the relation between attention effect in EEG power for the indicated frequency bands and laminar fMRI connectivity. The signed squared partial Spearman correlation over subject (n=30) is shown. For each frequency band, the effects of other frequency bands and pupil size are partialed out. The p-values to the largest supra-threshold cluster in each sub-panel and are based on a non-parametric cluster-based permutation test. Results for individual region combinations are depicted in supporting Figs. S1-S4. The relation between pupil size and inter-regional connectivity is depicted in Fig. S5. The results for inter-regional connectivity based on the top 5% and top 25% activated vertices are shown in supporting Figs. S6 & S7.

**Fig. 4:**
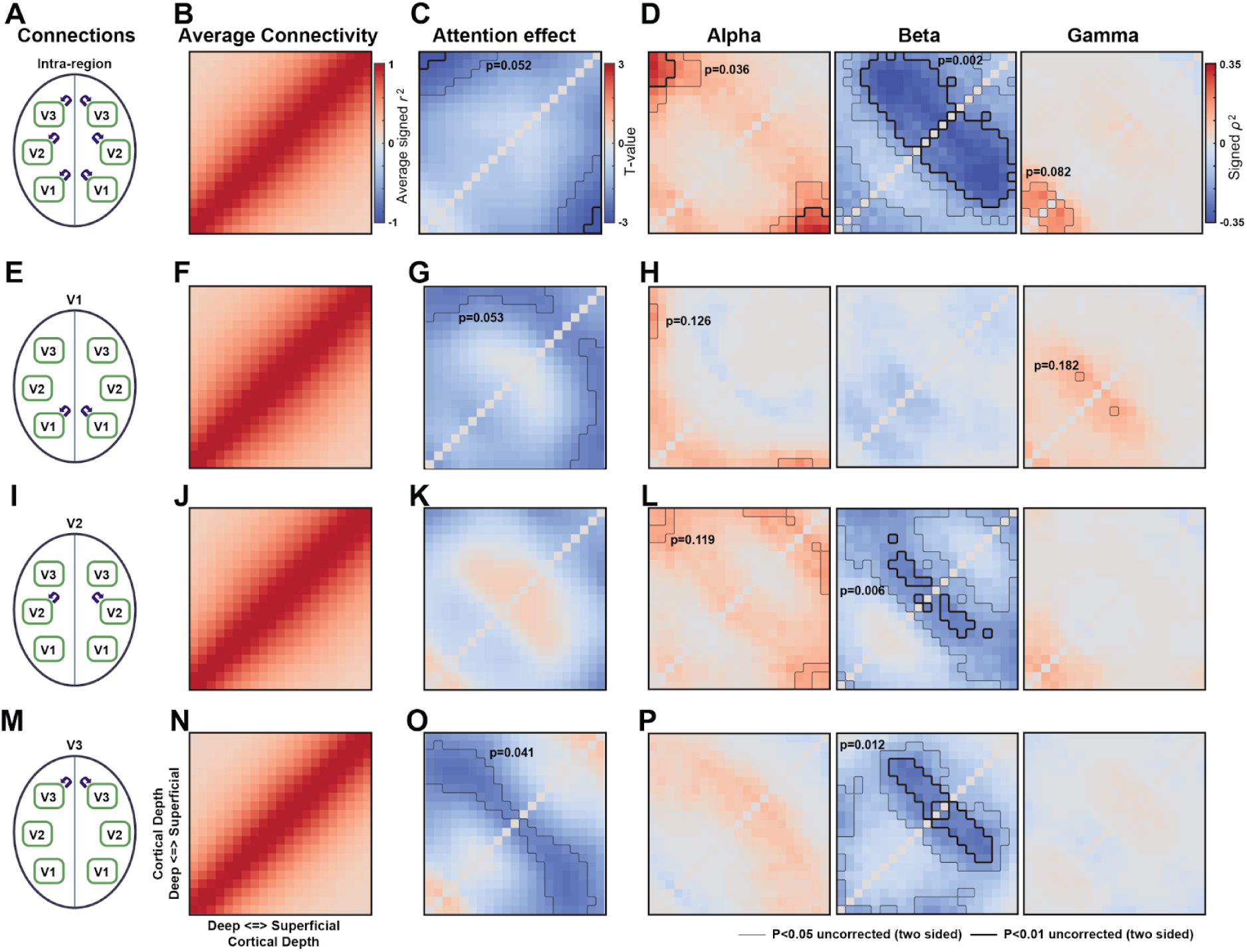
Relation between EEG power and intra-regional laminar fMRI connectivity. The organization of the columns, rows, panels and sub-panels and the information they contain is the similar as for Fig. 3. Since connectivity within regions is depicted here, the same region is here plotted on the y-axis and the x-axis. As a consequence, the results are symmetric about the x=y diagonal. Results for individual regions are depicted in supporting Figs. S1-S4. The relation between pupil size and intra-regional connectivity is depicted in Fig. S5. The results for intra-regional connectivity based on the top 5% and top 25% activated vertices are shown in supporting Figs. S6 & S7.

A schematic representation of how we computed laminar connectivity and its relation to EEG power is depicted in Fig. 2. In panel A we illustrate how we compute the attention effect in laminar connectivity for a single inter-/intraregional combination of laminar depths, which was repeated for all region/depth combinations. We computed the attention effect in laminar connectivity in the following steps: First, we estimated the amplitude of single trial BOLD responses for the two conditions without a speed-increase (attention ON and attention OFF) by fitting a canonical hemodynamic response function. Subsequently, for each condition separately, the correlation of these amplitudes over trials was calculated (excluding erroneous trials and trials with eye blinks). After applying a Fisher z-transformation (i.e. arctanh(r)) to account for ceiling effects, these were averaged over the separate task blocks and grouped into interregional connectivity within a hemisphere, and intraregional connectivity within the brain regions (see panels A of Fig. 3 and Fig. 4). Subsequently for each of the groups the attention contrast was calculated by subtracting the connectivity-metric for the “attention OFF” condition from the “attention ON” condition, the result of this analysis is illustrated in panel B. In order to relate EEG frequency bands to laminar fMRI connectivity, this attention contrast was then correlated over subjects (panel D) separately with the attention effects in respectively the alpha, beta and gamma bands (panel C) using a partial Spearman correlation (correcting for effects in the other frequency bands and in pupil size (Fig. S1)). This results in a two-dimensional representation of how laminar fMRI connectivity relates to a specific EEG frequency band, illustrated in panel E. The results of these analyses are depicted in panels B-D of Fig. 3 and Fig. 4.

In Fig. 3 & Fig. 4, the first column shows which connections are being taken into account to obtain the results in each row. The top row being the average for all possible combinations, the other rows showing the results for individual region combinations where the only averaging that was applied was over hemispheres. Throughout this paper we will primarily focus on the results of the top row, i.e. the average over all regions. The supporting figures contain the results for each region separately and their correlation with EEG (supporting Figs. S1-S4). In the original analysis of this data, we observed an attention effect in pupil size, which is depicted in Fig. S5, where we also show the correlation of the pupil-attention effect with laminar connectivity within and between regions, observing no significant effects

Fig. 3B depicts the average fMRI connectivity summed over the two conditions for inter-region connectivity while Fig. 4B shows the same for intra-regional connectivity. Inter-region connectivity increased along the diagonal from deep-to-deep connectivity pairings to superficial-to-superficial pairings, in accordance with previous observations (Wu et al 2018). Due to partial volume effects, connectivity is strongest close to the diagonal for intraregional connections (on the diagonal it is 1 by definition). In panel C of Fig. 3 & Fig. 4 we depict the attention effects in BOLD connectivity. Between regions (Fig. 3), no strong laminar specific effects were observed, although there is a global trend that indicates a relative decrease with attention. Within brain regions (Fig. 4), a relatively strong, negative effect was found for connectivity between deep and superficial layers, as well as a non-specific decrease with attention similar to the one observed in Fig. 4C.

In panel D of Fig. 3 & Fig. 4. we show how the attention contrast in laminar connectivity correlates with the attention effect in EEG power. The most prominent observation is that decreased beta band power strongly relates (over subjects) to increased laminar specific connectivity interregionally as well as intraregionally. For interregional laminar interactions, beta strongly correlates over subjects with deep-to-deep layer connectivity, while for intraregional connectivity it relates to deep/middle to middle/superficial connectivity. The negative correlation with beta power is also observed for laminar connectivity between individual region pairs (Fig. S3). This indicates that the observed task related decrease in beta band power relates to both increased intra- and interregional laminar fMRI connectivity. For the alpha band we observe a *positive* correlation with intraregional deep-to-superficial layer connectivity. Although there is no strong effect observed for alpha for interregional connectivity, the trend seems to be the same, as well as for the relation of alpha with connectivity between single pairs of brain regions (Fig. S2). Contrary to the beta effect, the attention related alpha power decrease seems therefore related to a decrease in connectivity. The relation of EEG alpha power with deep-to-superficial intraregional connectivity shows a similar pattern to the attentional decrease in fMRI connectivity (compare Fig. 4C to D). To verify whether the effects observed for the alpha and beta bands are stable over analysis strategies, we repeated the analysis for the laminar signals extracted from the top 5% (Fig. S6) and top 25% (Fig. S7) activated signals. These analyses yielded qualitatively similar results as those presented here.

The results for the individual region-pairs suggest that inter- and intraregional connectivity are in general consistent with the average over individual region combinations, but also have unique features. In line with the strong negative correlation between beta power and deep-to-deep layer coupling, for all three individual region pairs (V1-V2, V1-V3, and V2-V3) a negative correlation is observed for deep-to-deep layer coupling. Similar patterns can also be observed for correlations of alpha and beta power with intra-regional connectivity (averaged over all regions) and correlations with connectivity within V1, V2 and V3 in isolation. The effects for the individual regions seem here in general weaker as for the combined results for intraregional connectivity.

Besides these general patterns, the results for the regions-pairs and regions in isolation also reveal some unique features. In particular, the relation between beta and interregional coupling shows variation over the individual region-pairs where the V1-V2 and V2-V3 also contain unique features: For V2-V3, an additional peak for beta power is observed for the coupling between deep layer V3 and both deep and superficial layers in V2. Similarly, for V1-V2, additional beta peak is present between deep V1 and middle layer V2.

## DISCUSSION

In this study, we explore how task related changes in fMRI connectivity within and between brain regions relate to changes in neural synchronization measured by frequency specific EEG power. We used an attention demanding visual change detection task, which allowed us to correlate task induced differences in alpha, beta and gamma power to depth dependent connectivity between regions and within regions. We observed the strongest link between laminar fMRI connectivity and EEG power for the beta band. The negative correlation between beta power and deep-to-deep interregional layer connectivity as well as deep/middle to superficial layer connectivity indicates that a decrease in beta power reflects an increase in laminar specific connectivity. For the alpha band we observe a positive correlation pattern for intraregional connectivity which strongly resembles the inverse of the pattern of decreased fMRI connectivity as a result of the attentional modulation. For both the attentional modulation as well as the correlation with alpha power, a strong effect was observed for the deep-to-superficial intraregional layer connection. For interregional connectivity no significant relation with the alpha band was observed, although in general, a positive relation was present as well. This indicates that, in contrast to beta band decrease, the task induced alpha band decrease relates to decreased laminar connectivity, particularly within a region. For the gamma band, no relation was observed with laminar connectivity within regions, indicating that while gamma band activity is related to the strength of the BOLD signal in middle and superficial layers, this is not reflected in changes in laminar fMRI connectivity within and between brain regions. The general patterns observed for the aggregated inter- and intra-regional coupling were also reflected in the individual region pairs (inter-regional) and regions (intra-regional), although unique features were also observed, in particular for the relation of beta power with coupling between V1-V2 and V2-V3.

The results for the beta band presented here support a notion that especially deep-to-deep layer feedback projections are related to beta band activity (Bastos et al 2012, Bastos et al 2015, Buffalo et al 2011). This is further supported by our previous finding from this dataset that beta power correlates negatively with BOLD activity in deep layers (Scheeringa et al 2016), which aligns well with observations in EEG and MEG that beta power, like alpha power, often decreases in regions that become actively involved in a task (Spitzer & Haegens 2017). On the other hand, inter-areal beta-band synchrony is regularly reported to increase when brain regions become task involved (Bastos et al 2015, Buschman & Miller 2007, Schoffelen et al 2017). This leads to a pattern of locally decreased beta band activity, reflected in a power decrease in EEG and MEG, and greater inter-regional beta band synchronization, which is typically better measured intracranially in animals. Here we find the deep-layer inter-regional connectivity increases for subjects that show a stronger task related beta power decrease. Following from the pattern described above, this increase might reflect the same underlying process that is reflected in (deep-to-deep layer) interregional beta band synchrony observed in animals. In addition, within a brain region we observed that a decrease in beta power predicts increased connectivity between deep/middle and superficial layers. Combined with the observation that beta correlates negatively with deep layer BOLD, and deep-to-deep layer interregional connectivity, these findings suggest that within a brain region a decrease in beta power reflects increased incoming feedback related information in deep layers that is subsequently projected to middle/superficial layers.

For the alpha band, our main finding is that decreased alpha power (in the attention ON compared to the attention OFF condition) relates strongly to decreased correlations between deep and superficial layer BOLD-responses within the same region. This is in line with the observation in laminar recordings in monkeys that connected alpha sources can be found in both infragranular and supragranular layers (Bollimunta et al 2008, Bollimunta et al 2011, Haegens et al 2015). Typically, alpha is seen as either an idling rhythm that occurs when the cortex is not task involved, or a reflection of top-down inhibition of task irrelevant brain regions (Jensen & Mazaheri 2010, Klimesch et al 2007, Pfurtscheller et al 1996a). The results here suggest these processes might be reflected in greater correlations between deep and superficial layer activity within regions. The inhibition or idling function of an increased alpha amplitude seems to be reflected in lower, but more correlated, neural activity across layers. This increased correlation might reflect a lower capacity for independent and differentiated neural responses in different layers of the cortex. Task evoked decreases in inter-neural coupling have been previously observed in in spike rate correlations and fMRI connectivity (Cohen & Maunsell 2009, Cole et al 2014, Ecker et al 2010, Gonzalez-Castillo & Bandettini 2018, Ito et al 2020, Ruff & Cohen 2014). Our results here suggest these decreases relate to task-evoked changes in local neural synchronization in the alpha band.

A noteworthy observation here is the lack of a relation between gamma band activity and laminar connectivity. Gamma band activity is thought to reflect feed-forward flow of information from lower order superficial to higher order middle cortical layers(Bastos et al 2012, Bastos et al 2015, Bosman et al 2012, Fries 2015) which is reflected in increased coupling between (laminar resolved) electrophysiological measures from different regions (Bosman et al 2012, Grothe et al 2012, Michalareas et al 2016, Roberts et al 2013, van Kerkoerle et al 2014). In line with this, we found gamma band activity to correlate positively with middle/superficial layer BOLD responses to visual stimulation (Scheeringa et al 2016). We however did not find this reflected in a relation between gamma and laminar connectivity. Our results therefore suggest that neural and cognitive processes related to laminar specific interactions with increased gamma-band synchrony might not be well studied by approaches based on fMRI-based laminar-level coupling.

Our observation that alpha and beta band activity have relationships with fMRI laminar *connectivity* that differ in sign (especially within a brain region) is surprising, since both bands show an attentional decrease in power and negatively correlate with the BOLD signal (Fig. 1D). Since alpha is thought to indicate functional inhibition or idling of task irrelevant regions (Jensen & Mazaheri 2010, Klimesch et al 2007, Pfurtscheller et al 1996a), our results suggest increased BOLD connectivity, most pronounced for connectivity between layers within a region, can be a sign of decreased task involvement of connected but separate neural populations. Likewise, beta power is often observed to decrease in a context where regions become involved in task execution and increase when involvement stops, as is illustrated by the motor related beta-rebound (Pfurtscheller et al 1996b, Salmelin et al 1995). However, for beta, higher connectivity within and between regions is observed for subjects that showed a larger task related power decrease. Combined, these results for the alpha and beta bands indicate that an increase as well as a decrease in (laminar) fMRI connectivity measures can indicate greater involvement in the same task of separate neural populations (e.g. in different layers or regions). In general, this observation complicates the interpretation of fMRI based laminar and non-laminar connectivity on their own. Being able to relate (laminar) fMRI connectivity effects to EEG measures can facilitate this as is demonstrated here. However, a comparison of the pattern of attention effects in connectivity between regions in Fig. 2C with the observed relation with EEG frequency band in Fig. 2D, suggests that the pattern in attention effects cannot fully be explained by the EEG frequency bands under investigation here. This indicates that there are influences on task related fMRI connectivity changes that are not reflected in EEG power. It is important to note that fMRI does not measure neural activity in the same way as EEG and MEG. The latter critically depend on phase-synchronous responses in post-synaptic membrane potential of a part of the underlying neural population (apical dendrites of pyramidal cells), while fMRI can better be considered a representation of, temporally low-pass filtered, total synaptic activity of the measured population.

Combined with the observation that task effects in alpha correlated with task effects in superficial layer BOLD and beta with task effects in deep layer BOLD, the laminar fMRI results indicate distinct neural mechanisms underlying the alpha and beta effects, and therefore likely relate to distinct cognitive processes. In experiments that test the predictive coding framework, and also in animal electrophysiological work in general, these frequency bands are often not assessed separately (Bastos et al 2020, Bauer et al 2014, Buffalo et al 2011). The finding that stronger alpha oscillations relate to stronger coupling between neural populations, might reflect a less differentiated neural response to stimuli. Within a predictive coding framework, this increased coupling might be interpreted with decreased precision (of the prediction) when alpha amplitude increases, while the predictions themselves are related to beta band oscillations.

EEG power is believed to be a reflection of synchrony between neurons in post-synaptic potentials within a specific frequency band over a larger part of the cortex (Lopes da Silva 2013). Consequently, stronger coupling between neurons is expected when synchrony (e.g. EEG power) is higher regardless of the frequency band. Therefore, a positive correlation could be expected for all three frequency bands investigated here. We do not find this. Only for alpha power do we find stronger coupling between layers within regions. For beta power we find the opposite effect both within and between regions, while for gamma power no relation was observed for either. There are several factors that could be related to this. The neural synchrony we measure with EEG is related to millisecond level synchronization of post-synaptic potentials of only a sub-population of neurons (parallel oriented pyramidal cells). The fMRI BOLD signal is more related to the total amount of peri-synaptic activity as far as it results in a local change in blood flow, and therefore does not depend on the millisecond level synchrony necessary for detectable oscillations in EEG, and is also the direct consequence of neural activity in a larger neural population. The relation between frequency specific EEG synchrony and laminar connectivity therefore is also likely a relation between at least in part different types of neural activity.

The ability to study fMRI connectivity at the laminar level is a relatively recent development. Besides linking laminar the fMRI connectivity to frequency specific EEG power, the findings here also provide insight to laminar fMRI itself. A first pattern that emerges from our analysis is that irrespective of attention condition, we observed that connectivity increased along the diagonal from deep-to-deep connectivity pairings to superficial-to-superficial pairings. This is in line with earlier observations of connectivity between voxels within a region (Wu et al 2018), and is likely related to the larger contribution to the BOLD signal from draining veins which increases towards the surface of the cortex. When we subsequently compare the attention ON versus the attention OFF condition, the outcomes do suggest a general decrease in fMRI connectivity with attention for both within and between region connectivity, although no significant results were observed for our crude attention modulation in laminar fMRI connectivity within and between regions. While no particular laminar pattern stands out for connectivity between regions, for within region connectivity a pattern of decreased coupling between deep and superficial layers stood out that resembles the positive correlation observed with alpha power (Fig. 4, panels C & D). Since our task manipulation is not a clean modulation of attention but also includes cognitive processes like arousal and the predictability of the speed change, these decreases in connectivity cannot directly be linked to a specific cognitive process. It is however not uncommon to observe task-evoked decreases in neural correlations within a region and between connected cortical regions in both neural spiking and fMRI connectivity (Cohen & Maunsell 2009, Cole et al 2014, Gonzalez-Castillo & Bandettini 2018, Ito et al 2020, Ruff & Cohen 2014). Since our experimental manipulation essentially consisted of a task ON versus task OFF contrast, our results might reflect the same or a similar process. Our results suggest that this process of decorrelation of neural populations, especially between different layers in the same region, is related to a decrease in alpha power.

The results obtained for intra- and interregional connectivity are aggregated over individual region combinations. The pattern of the relation between EEG power and laminar connectivity we observe for the aggregates is in general in line with the individual laminar connectivity patterns, suggesting the aggregate results reflect a more general pattern. These individual laminar connectivity patterns however also contain some unique features, in particular for the relation of beta power with connectivity between V1 and V3, and between V2 and V3. For V2-V3, beta power is found to relate to the coupling of deep layer V3 to both deep and superficial layers in V2, a pattern that might reflect anatomical feedback projections that go from deep layer V3 to both deep and superficial layers in V2. The stronger correlation of beta with coupling between deep V1 and more middle V2 is however harder to interpret since direct anatomical connections are largely absent here. In general, we want to caution against over-interpreting all the different patterns yielded by analyses. In our view, patterns that recur over the different analyses (over individual region combinations and different approaches of region/voxel/vertices of interest selection) are more likely to reflect reliable true patterns, than variations in specific sub-analyses. The analyses in this study are based on thirty subjects, making it one of the larger studies using laminar fMRI. It is however still on the small side compared to other studies relating individual differences in (f)MRI based measures to other measures over subjects. Therefore, we see this study more as a proof of principle that demonstrates that changes in fMRI based laminar connectivity relate to changes in the strength of EEG connectivity. The exact pattern and nature will no doubt be subject of further research.

In conclusion, this exploratory study provides the first evidence that neural oscillations measured by EEG reflect not only laminar specific neural activity, but also changes in laminar specific connectivity within and between brain regions. As such, it provides a neurophysiological basis for investigating laminar level connectivity with fMRI. It further suggests that the neural processes underlying alpha and beta oscillations are associated with opposite effects in correlation based connectivity measures in (laminar) fMRI, while their relations to the strength of the BOLD response and modulation by task conditions are highly similar. If this observation is indeed further verified, this could have important implications for correlation based measures of functional and effective connectivity. This would imply that it is crucial to understand in which conditions alpha and beta neural oscillations are modulated in order to understand the effects of (laminar) connectivity changes in fMRI. Integrated analysis of laminar fMRI connectivity with electrophysiological measures like EEG and MEG might provide further insights on this topic.

## Acknowledgements

This work was funded by the Emmy Noether Programme of the Deutsche Forschungsgemeinschaft (DFG), Grant KO 5341/1-1 to Peter Koopmans. René Scheeringa and Mathilde Bonnefond acknowledge support for the European Research Council under the European Union’s Seventh Framework Programme (FP7/2007–2013)/ERC starting grant agreement no 716862. The data was recorded in a project funded by a grant from The Netherlands Organization for Scientific (NWO) to René Scheeringa (Veni scheme 451-12-021).

## METHODS

### Relevant methods from previous work

The work presented here builds upon the data recorded and analyses performed in our previous study (Scheeringa et al., 2016). The parts of the methods in that study that are relevant for the novel analysis presented here were taken from the methods described in the supporting information for this publication. The experimental paradigm and parts of the methodology this analysis and our previous work was based on were adapted from a previous older experiment described in Scheeringa et al. (Scheeringa et al 2011) and are in some parts very similar. Parts of the section below are therefore indirectly taken from that article as well and adapted to the current analyses.

#### Subjects

In this article we analyzed the data of the same thirty subjects that were included in the analysis of our earlier work and this section is in parts the same and in parts adapted from the same section in Scheeringa et al. *(2016)*.

In this study we a priori chose to include 30 subjects. To achieve this we measured, 34 subjects (29 female, 5 male, mean age 21.6 y, range 18–26 y) without a history of known psychiatric or neurological disorders participated in the simultaneous EEG/fMRI session. All had normal or corrected-to-normal vision. Before the start of the experiment, written informed consent was obtained from each subject. The experiment was approved by a local ethical committee (CMO Arnhem/Nijmegen region, protocol 2001/095). Subjects for whom a sustained gamma-band response to visual stimulation was observed (see EEG Data) were selected for further analysis. After this selection, 30 subjects (26 female, 4 male) remained. All of the results presented here are based on these 30 subjects. Subjects were paid a small fee (€35, and €5 for each half hour the experiment took over 3.5 h) or given course credits. Subjects were recruited via an online system for subject recruitment at the Radboud University Nijmegen and were mainly recruited from the student population. Subjects with a relatively small head size were requested (≤58 cm circumference) because pilots indicated that large heads would not fit within with the relatively small field of view of the high-resolution functional scans. This decision also contributed to the relatively large proportion of female participants.

#### General Experimental Procedure

Subjects came to the Donders Institute in Nijmegen 1 h and 15 min before the start of the experiment. During this time, the procedures and task were explained to them, written informed consent was obtained, the EEG electrodes were applied, and the electrode positions were recorded using a Polhemus FASTRAK. After these steps, the subjects were positioned in the MRI scanner for simultaneous EEG–fMRI registration. The head was stabilized with foam pillows underneath and at the side of the head, and a strand of paper adhesive tape was applied to the forehead to provide feedback to minimize head movements. The MRI-compatible EEG amplifiers were positioned at the right foot end of the subject. When in the scanner, first a normal (1 mm) resolution T1 anatomical scan was obtained. During this scan, subjects performed a practice version of the main task to familiarize themselves with it. This practice version was followed by a shorter blocked version of the task that allowed us to run a simple GLM analysis on the MR console, the results of which were used as a localizer in the positioning of further scans. The slices of the high-resolution (0.75 mm) 3D EPI scans were planned to coincide with the main activations in the early visual cortex, and a few test volumes were acquired, while avoiding the eyes. Subsequently, a high-resolution (0.75 mm) anatomical T1 scan and a field map were made using the same slice orientations as for the high-resolution EPI volumes. This step was followed by three main task blocks, each with a duration of roughly 20 min, during which simultaneous EEG and high-resolution fMRI data were acquired. In between blocks, subjects were allowed to rest for a few minutes. After the third block, another field map was acquired, followed by a short retinotopy measurement using a normal 2D EPI sequence. In total, subjects were in the MRI scanner for ∼1 h and 45 min.

#### Experimental Paradigms

##### Visual attention task

Subjects engaged in a visual attention task that is known to elicit strong, long-lasting (up to several seconds), and narrow-band gamma-activity increases, as well as alpha- and beta-band decreases in both MEG (Fries et al 2008, Hoogenboom et al 2006, Marshall et al 2015, Muthukumaraswamy et al 2009, Muthukumaraswamy & Singh 2008, Muthukumaraswamy & Singh 2009) and EEG (Fries et al 2008, Koch et al 2009, Scheeringa et al 2011). Source analysis has revealed that these frequency-specific responses originate from the early visual cortex(Fries et al 2008, Hoogenboom et al 2006, Marshall et al 2015, Muthukumaraswamy et al 2009, Muthukumaraswamy & Singh 2009) and overlap with task-evoked fMRI activations (Fries et al 2008)(28). In this task, subjects attended to circular, inward moving gratings, and were asked to detect a change in inward speed.

Each trial started with a reduction in contrast of a fixation point that was present between trials (Gaussian of 0.4°) by 40%. This contrast reduction served as a warning for the upcoming visual stimulation and instructed the subjects to stop blinking until the end of the trial. After 1,600 ms, a red “!” or a green “=“ appeared just above the fixation point for 100 ms. The “!” indicated a 66.7% chance of an increase in the speed of the upcoming inward moving grating and was presented in 75% of the trials. The “=“ indicated that no speed change would occur with 100% accuracy and was presented in 25% of the trials. After this attention cue was removed, the fixation point remained on the screen for 400 ms before it was replaced by a sine wave grating (diameter, 7°; spatial frequency, 2.5 cycles per degree; contrast, 100%). The sine wave grating contracted to the fixation point (1.6° per second) for one of four stimulus durations: 1,200, 1,400, or 1,600 ms. This step was followed by an increase in the contraction speed to 2.2° per second for maximally 500 ms after either 1,200 or 1,400 ms. The trials with 1,600-ms stimulation were not followed by a speed change. Therefore, trials cued with “=“ were always followed by a 1,600-ms visual stimulation period whereas, for trials cued with “!”, this happened in 33.3% of the trials. An equal number of trials of the four conditions were presented (1,200-ms stimulation, attended, speed change; 1,400-ms stimulation, attended, speed change; 1,600-ms stimulation, attended, no speed change; 1,600-ms stimulation, not attended, no speed change).

Subjects were instructed to press a button with their right index finger as soon as they detected the speed change. The stimulus disappeared after a response was given, after 1,600 ms of stimulation (for catch trials), or, if no response was given, within 500 ms of the speed change. Feedback about the performance was given for 500 ms. In the case of a correct response or if a response was correctly withheld, “ok!” appeared in green above the fixation point. In case of premature or slow/no responses, “early” or “late,” respectively, appeared in red.

Subjects performed a practice version of 40 trials of the task during the low-resolution anatomical scan at the start of the session. This practice block was 5-min, 28-s long, and a trial was presented every 8.2 s. Subsequently, a localizer version of this paradigm was performed to obtain an activation map that was used to plan the slices for the orientation of the slices of the high-resolution fMRI volumes (see MRI Data Acquisition). To perform a simple activation versus “rest” analysis, 64 trials were grouped in 8 blocks of 32 s, with 8 trials each, and a trial starting every 4 s within a block of 8 trials. Each group of eight trials consisted of two trials of each condition. Within a group of eight trials, the order was randomized. This block was followed by three blocks during which EEG was combined with high-resolution fMRI (i.e., the “main task”). Each session consisted of 72 trials, 18 of each condition. Onset of a trial was triggered by every third MRI volume. The laminar resolution MRI sequence we used consisted of the acquisition of three volumes of 3.792 s, followed by a scan-free period of the same length to allow for MR gradient and RF pulse-free recording of EEG data during the presentation of the trial. The trial, starting with the dimming of the fixation point, started 330 ms before the end of the third scan. The attention cue was presented in the artifact-free period, 1,260 ms after the end of the third scan, and stimulation onset was at 1,760 ms. A trial was presented every 15.168 s. In total, each session was 18-min, 12-s long.

##### Retinotopy

To separate early visual regions (V1, V2, and V3), a clock-wise rotating double (one in each hemifield) wedge-shaped red-green flickering checkerboard was used. The flicker frequency was 8 Hz. The two wedges started at the center of the visual field, had an angle of 22.5°, and were presented in opposition: i.e., at a 180° angle from each other. By simultaneously using wedges in the left and right visual fields, we were able to simultaneously map left and right early visual regions. The wedges remained stationary for 6 s, after which they skipped to the next position in a clockwise fashion. After eight displacements, all angles of the visual field were stimulated. This sequence was then repeated eight times, which resulted in a duration of 6 min and 24 s for the retinotopy measurement.

#### MRI Data Acquisition

MRI data were acquired using a 3.0-T whole body MRI scanner (Magnetom Trio Tim; Siemens). A 32-channel head-coil was used to record the images. For the 1-mm isotropic T1-weighted anatomical scan, a 3D magnetization-prepared rapid gradient-echo (MPRAGE) sequence was used [inversion time (TI), 1,100 ms; repetition time (TR), 2,300 ms; 192 slices perslab; 1-mm slice thickness; generalized autocalibrating partially parallel acquisition (GRAPPA) acceleration factor, 4; field of view (FOV), 256 mm; voxel size, 1.0 by 1.0 by 1.0 mm], which took 3 min and 8 s to acquire. This anatomical scan had whole brain coverage and was used to plan the slices of the high-resolution functional scans, in combination with the results of a functional localizer scan that were overlaid on it. This overlay showed all regions that were activated by the task (threshold, t > 4.00). The acquisition of this functional localizer directly followed the acquisition of the anatomical scan. Sequence parameters were as follows: 2D EPI, 3-mm isotropic resolution; echo time (TE), 30 ms; TR, 2,000 ms; 30 slices with 10% gap, 64 matrix, and 256 volumes.

A higher resolution structural scan with limited coverage was acquired, which was later used to construct cortical surfaces that were used to assign voxels to cortical depth bins. This scan had an isotropic resolution of 0.75 mm and 80 slices, other sequence parameters being identical to the whole-brain protocol. The center of the acquisition slab was aligned with the center of the slab of the high-resolution functional scans.

The high-resolution functional scans were acquired using a 3D EPI sequence (Poser et al 2010). Sequence parameters were as follows: TE, 30 ms; TR, 79 ms; 18° flip angle; 0.75-mm slice thickness; GRAPPA acceleration factor, 4; FOV, 192 mm; voxel size, 0.75 mm isotropic. The vendor’s default fat suppression was used. During piloting, we found that data quality improved markedly when using a large bandwidth-time product of 12 for the excitation pulse. Forty-eight slices were acquired, resulting in a volume acquisition time of 3.792 s. Slices were planned over the main activated regions in the visual cortex, based on the online analysis of the localizer task. In total, 219 functional images were recorded in each session, of which the first 3 images were discarded. To be able to measure EEG signals with as little interference as possible from the scanner, the scans were paused every three volumes for 3.792 s (i.e., one volume of pause). During this time, the stimuli were presented, and the EEG responses were recorded. The peak BOLD response had about a 6-s delay with respect to the stimulus onset. The BOLD response related to the visual stimulation therefore reached its peak approximately at the end of the first/beginning of the second volume after the pause block. The irregular excitation pattern (48 × 3 excitations spaced by 79 ms, followed by a pause of 3.792 s before the next excitation) caused the pseudo steady state of the MR signal to be perturbed. The perturbation was very regular, however, and was modeled in the GLM with an exponential sawtooth pattern that was computed from the gray matter signal (see Statistical models).

The functional scans were preceded and followed by 3-min field map measurements. The scanner shim settings were fixed after the first field map to make sure its results remained valid throughout the session. The field maps were acquired to help us perform a geometric distortion correction on the data. During the analysis stage, however, other research in our group resulted in an image-based distortion correction method for laminar analyses with superior performance, which was used instead. This method is summarized below in *Laminar-Level fMRI Analysis*.

The task activated a large part of the visual cortex. To be able to discriminate V1, V2, and V3, retinotopy was performed. FMRI data were acquired using a standard 2D EPI sequence with 2-mm isotropic resolution. Sequence parameters were as follows: TE, 30 ms; TR, 1,500 ms; flip angle, 70°; 2.5-mm isotropic resolution;24 slices; 64 matrix; 260 volumes were acquired.

#### Laminar-Level fMRI Analysis

The laminar fMRI analysis was performed following the strategy outlined in an earlier laminar fMRI paper (Koopmans et al 2011), with the inclusion of a new method to correct geometric distortions. The high-resolution 0.75-mm structural MPRAGE data were processed in Freesurfer (Dale et al 1999, Fischl et al 1999) to obtain a white matter (WM)–gray matter (GM) boundary surface mesh and a GM– pial surface boundary.

These surface meshes were overlaid on the high-resolution functional data after coregistration of the anatomical and average functional volume with the spm_coreg function in SPM (Friston 1995). However, differences in geometric distortion resulted in suboptimal boundary assignments in the functional data. This result was addressed using an in-house implementation of Boundary Based Registration (BBR) (Greve & Fischl 2009) that was modified to work recursively on ever smaller scales (Van Mourik et al 2014). The initial WM–GM surface mesh was fed into the BBR algorithm that calculated linear transformations of the mesh with a cost function to maximize the contrast gradient through the boundary. This contrast gradient is maximal when all white matter is on one side and all gray matter on the other. The same transformation was applied to the other surface mesh. The mesh was then split into parts, and the process was repeated for each of these to get a more optimal local transformation. This process was repeated several times such that each partition received a registration based on the local distortions. The registration of all subjects was visually inspected. After the meshes were aligned with the functional data, sampling intervals were drawn perpendicular to the cortex from each vertex of a mesh to its corresponding vertex on the other mesh. By sampling the functional data along these lines, through-cortex profiles of the fMRI signal were created using nearest-neighbor sampling as motivated earlier in Koopmans et al. (Koopmans et al 2011). During this sampling step, the cortical depth assignment was corrected for the cortical folding pattern according to the Bok principle (Bok 1929, Kleinnijenhuis et al 2015, Waehnert et al 2014). The resulting laminar profiles consisted of 21 bins.

To determine functional areas, retinotopy and main task GLM results were projected on the surface mesh representations, and regions of interest (ROIs) were drawn assigning each of the activated nodes on the meshes to a particular functional area. For the task activation, we first selected the 10% most activated nodes within each region in each hemisphere in terms of t values, using a regressor that modeled the visual stimulation period (see Statistical models for a description of the design matrix). For each cortical depth, a t value was computed. These t values were averaged over the cortical depth for each node. For this average, the top five bins (i.e., 25%) were not used to minimize the effect of larger veins on top of the cortex. All of the profiles for each area were averaged (500–3,000, depending on area size) to form a single profile for that area for each time point. The resulting data for each area, 21 × 216 arrays (bins × time points), were fed into a GLM containing EEG-based regressors. Because taking the top 10% activated vertices is an arbitrary selection criterion, we also calculated these region of interest signals based on the top 5% and top 25% activated vertices.

#### EEG Acquisition

EEG data were recorded with a custom-made MRI-compatible cap equipped with copper wired Ag/AgCL electrodes, including current limiting resistors (Easycap). Data were recorded from 63 scalp sites organized according to the international 10–20 system. One dedicated electrode was placed on the sternum to record the ECG. The reference electrode was placed at FCz. A 250-Hz low-pass analog hardware filter was placed between the electrode cap and the two MRI-compatible EEG amplifiers (BrainAmp MR plus; Brainproducts). The EEG was recorded with a 10-s time constant and continuously sampled at 5 kHz. EEG recordings were performed with Brain Vision Recorder software (Brainproducts GmbH).

#### EEG Data

##### Preprocessing

Downsampling of the EEG data were carried out in Vision Analyzer (Brainproducts GmbH). Further preprocessing was carried out in Fieldtrip (70), in which the data were first rereferenced to the common average. Subsequently, the data were segmented in epochs that started 1,300 ms before onset of visual stimulation and ended 400 ms after visual stimulation. These segments were visually checked for artifacts, and trials with anomalies such as eye blinks and large muscle artifacts were removed. Behaviorally erroneous trials were excluded from further analysis.

##### ICA-based denoising

To denoise the data, we used a two-step independent component analysis (ICA) approach adapted from a similar approach proposed by Debener et al. (Debener et al 2006). Each ICA step was performed with the Fast-ICA algorithm (Hyvarinen 1999).

In a first step, ICA was performed on the 50- to 90-Hz bandpass-filtered concatenated EEG-trials. The time period for estimation of the independent components started 300 ms before the cue and ended at the end of the stimulation period. The unmixing matrix thus obtained was applied to the unfiltered data. The resulting component time courses therefore had a broadband spectral content and were subsequently subjected to a time-frequency analysis as described in the previous section.

For the vast majority of the subjects, we were able to observe a sustained gamma-band response in some of the independent component time courses, with some interindividual variation in the peak frequency of this response. In a second step, we again applied ICA, but this time on EEG data that were bandpass-filtered with a narrow band of 20 or 25 Hz (depending on the spectral extent of each subject’s gamma-response) around the individually adjusted peak of the sustained gamma-response. The unmixing weights of this analysis were again applied on the unfiltered data, and a time-frequency analysis was run on the component time courses.

Components that showed a sustained gamma-band response were selected (one to four components for each subject) and projected back to channel level. Finally, these channel-level data were again subjected to a time-frequency analysis, separately for the lower and the higher frequency windows, as described in the previous section. The results of this analysis constitute the basis for the construction of regressors that were used in the integrated EEG– fMRI analysis. With this strategy, we were able to denoise the EEG data across the different frequency bands that have been reported before using the same or similar paradigm (Hoogenboom et al 2006, Koch et al 2009, Muthukumaraswamy & Singh 2008, Scheeringa et al 2011). In total, we were able to observe these expected effects in 30 of the 34 subjects. This strategy based on gamma-band ICA proved to be the best strategy for denoising the EEG data across the different frequency bands, as has been previously reported (Scheeringa et al 2011).

##### Time-frequency analysis

Time-frequency analyses of the EEG data were carried out in Fieldtrip using a multitaper approach(Mitra & Pesaran 1999). To optimize the trade-off between time and frequency resolution, we carried out separate analyses for a lower frequency window (2.5–30 Hz) and a higher frequency window (30–120 Hz). For the lower frequencies, the power was estimated for windows of 0.8 s length moved across the data in steps of 100 ms. The frequency resolution was 1.25 Hz, and the use of three tapers resulted in a spectral smoothing of ±2.5 Hz. For the higher frequencies, the power was estimated for windows of 0.4 s length moved across the data in steps of 100 ms. The frequency resolution was 2.5 Hz, and the use of seven tapers resulted in a spectral smoothing of ±10 Hz. For both high and low frequencies, power was estimated up until the end of the visual stimulation period.

### Methodology new in this study

#### Relevant previous results

After using an ICA denoising strategy (Debener et al 2006), our previous analysis confirmed strong power decreases from baseline in alpha and beta bands, and increases in the gamma bands. Crucial for the analyses presented here, in the last second of the trial, alpha and beta power was lower for the attention ON condition compared to the attention OFF condition while gamma power was higher in this condition (Fig. 1B). For the cortical depth resolved fMRI analysis, we first estimated the BOLD signal at 21 depths from the boundary between white and gray matter to the boundary between gray matter and CSF. Subsequently we averaged the BOLD signal for each bin over vertices within regions of interest in bilateral V1, V2, and V3. For the main analysis presented here we selected the top 10% activated vertices collapsed over cortical depth within these regions. Although arbitrary, a threshold is necessary to select the regions within V1, V2 and V3 that are involved in the task. For the three regions BOLD was observed to be higher in the attention ON condition across all cortical depths. Besides these attention effects in EEG power and BOLD, we also observed that the pupil size was larger during the last second of the trial during attention ON. In our previous analyses, we repeated the integrated EEG-fMRI analyses for the top 5% and 25% activated vertices to verify whether to the arbitrary choice of the threshold affected the results. For the analyses presented here we do this as well.

#### Estimating Laminar Connectivity

In this study we estimate task related laminar fMRI connectivity from single trial BOLD amplitude estimates for all region/cortical depth combinations of the six regions involved. This was carried out by fitting the canonical HRF as implemented in SPM 12 (https://www.fil.ion.ucl.ac.uk/spm/) for every trial of the attention OFF and attention ON conditions without an increase in contraction speed. The other two conditions were each modeled by one regressor. The other included confound regressors were (i) two regressors modeling the button presses in the two trial types with a speed change; (ii) two regressors modeling the reaction time as a parametric modulation (one for each trial type with a speed change); (iii) regressors modeling the behaviorally incorrect trials for the two conditions with a speed change; (iv) the realignment parameters, their squares and derivatives to control for possible movement artifacts; (v) a regressor modeling the T1 effect related to the pause after every third scan; and (vi) six sine and six cosine waves with frequencies up to 0.006 Hz. The regressor modeling the T1 effect was obtained by averaging the global gray matter signal over every first, second, and third scan after a trial separately, and appending these same three values 71 times to form a regressor of 216 data points in length. The gray matter mask was obtained using a unified segmentation approach (Ashburner & Friston 2005) of the high-resolution anatomical scan. As a measure for connectivity we subsequently correlated for each condition the beta/parameter estimates of for every layer region/layer combination in every block for every subject. For the computation of this correlation, erroneous trials and trials including eye blinks were excluded. Subsequently, a Fisher-z transformation was applied to the correlations to account for the ceiling effects in correlation values. These values were then averaged over the task blocks and the region-combinations within each connectivity grouping as described in the first columns (Panels A, E, I & M) of Fig. 3 and Fig. 4. The effect of the attention manipulation was calculated as the difference between the attention OFF and the attention ON estimate of laminar connectivity between these groupings. This attention effect was subsequently used to correlate over subjects with the attention effects in alpha, beta and gamma power and in pupil size.

#### Correlating laminar connectivity with EEG power and pupil size

We correlated the attention effects in the alpha (7.5–12.5 Hz), beta (21.25–27.5 Hz), and gamma (50– 80 Hz) bands as well as the attention effect we observed in pupil size with the attention effects in laminar connectivity for the four groupings. For these bands, we averaged over the attention effect, expressed as a log-ratio of the attention-on versus attention-off condition, over all time bins from 500 ms until 1,600 ms after the onset of visual stimulation and all frequency bins within the indicated ranges. For the pupil size, the log-ratio of the attention effect in pupil size was averaged over the same time-period. The partial Spearman correlation (assuming a monotonic, but not linear relationship), was calculated between the attention effects in laminar connectivity and the EEG frequency bands and pupil size. For each frequency in question, the effects in the other two frequency bands and pupil size were controlled for. For pupil size the correlation was controlled for the three EEG frequency bands.

#### Statistical testing

Significant attention effects in laminar connectivity were tested using a non-parametric cluster-based permutation test (Maris & Oostenveld 2007) implemented in Fieldtrip (Oostenveld et al 2011). This procedure effectively controls the type-1 error rate in a situation involving multiple comparisons. This procedure was also used to test for testing task related differences in our original work (Scheeringa et al 2016). The procedure allows for user-defined test statistics tailored to the effect of interest within the framework of a cluster-based randomization test. Here, clusters were defined by adjacent layer-by-layer combination for which p<0.05 for an uncorrected two-tailed t-test. Within each cluster, the sum of all t-values were computed. These were compared with a reference distribution of cluster sums that was estimated from 200 000 permutations where the condition labels for each subject were randomly assigned. For each permutation the maximum cluster-sum was entered in the reference distribution.

Significant correlations of attention effects in EEG power and pupil size with attention effects in laminar connectivity were determined by a non-parametric cluster based approach (Maris & Oostenveld 2007) applied to correlations. After removing attention effects in other frequency bands/pupil size than the frequency band under investigation (effectively calculating a partial correlation), clusters were defined as adjacent layer-layer combinations for which the correlation passed an uncorrected significance threshold of p<0.05 (two tailed). The correlations were then transformed to t-values before they were summed within a cluster. These were compared with a reference distribution of cluster sums that was estimated from 200 000 permutations where the pairing between attention effects in EEG power and laminar fMRI connectivity was randomized over subjects. For each permutation the maximum cluster-sum was entered in the reference distribution.

