## Supporting Information for "Relating neural oscillations to laminar fMRI connectivity"

### Supplemental information

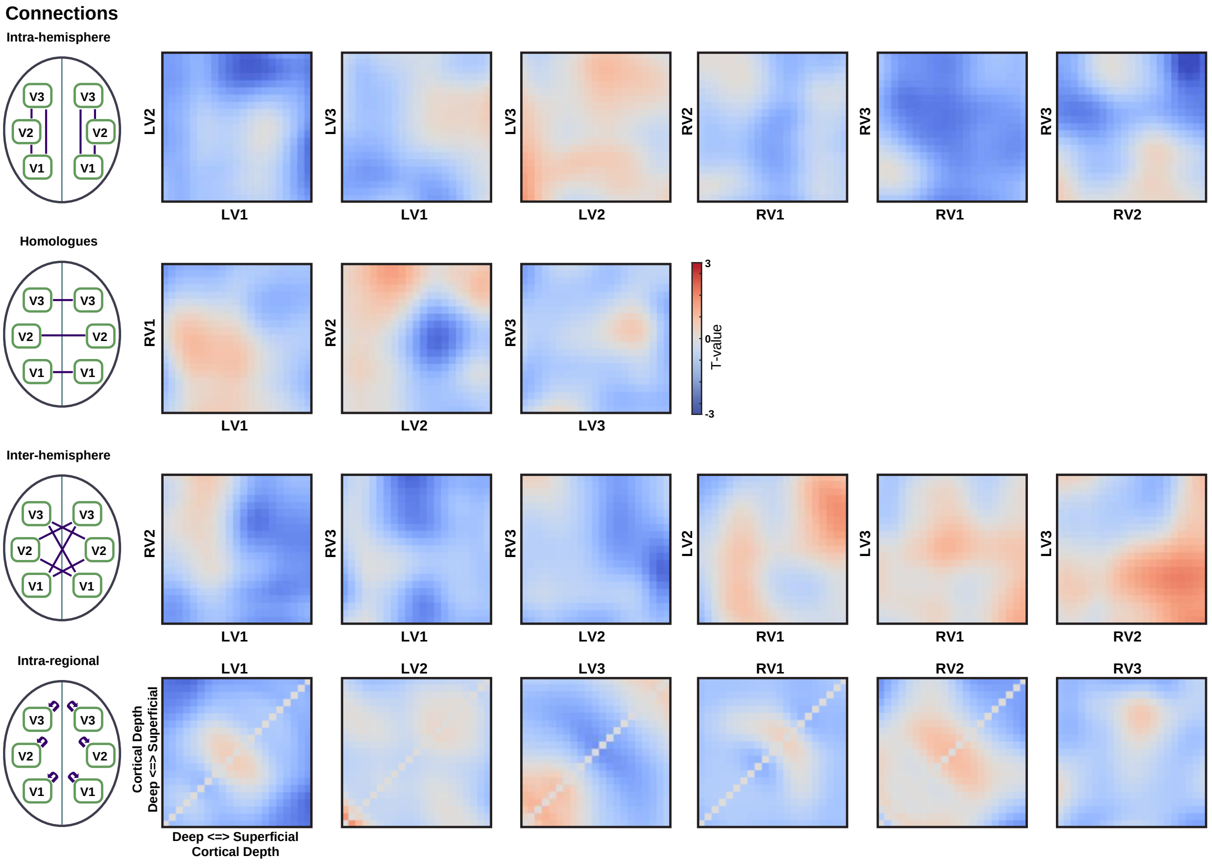


*Fig. S1: Attention effects for individual region pairs expressed in t-values, related to Figs. 3 and 4. Regions combinations are grouped as indicated in the left column.*

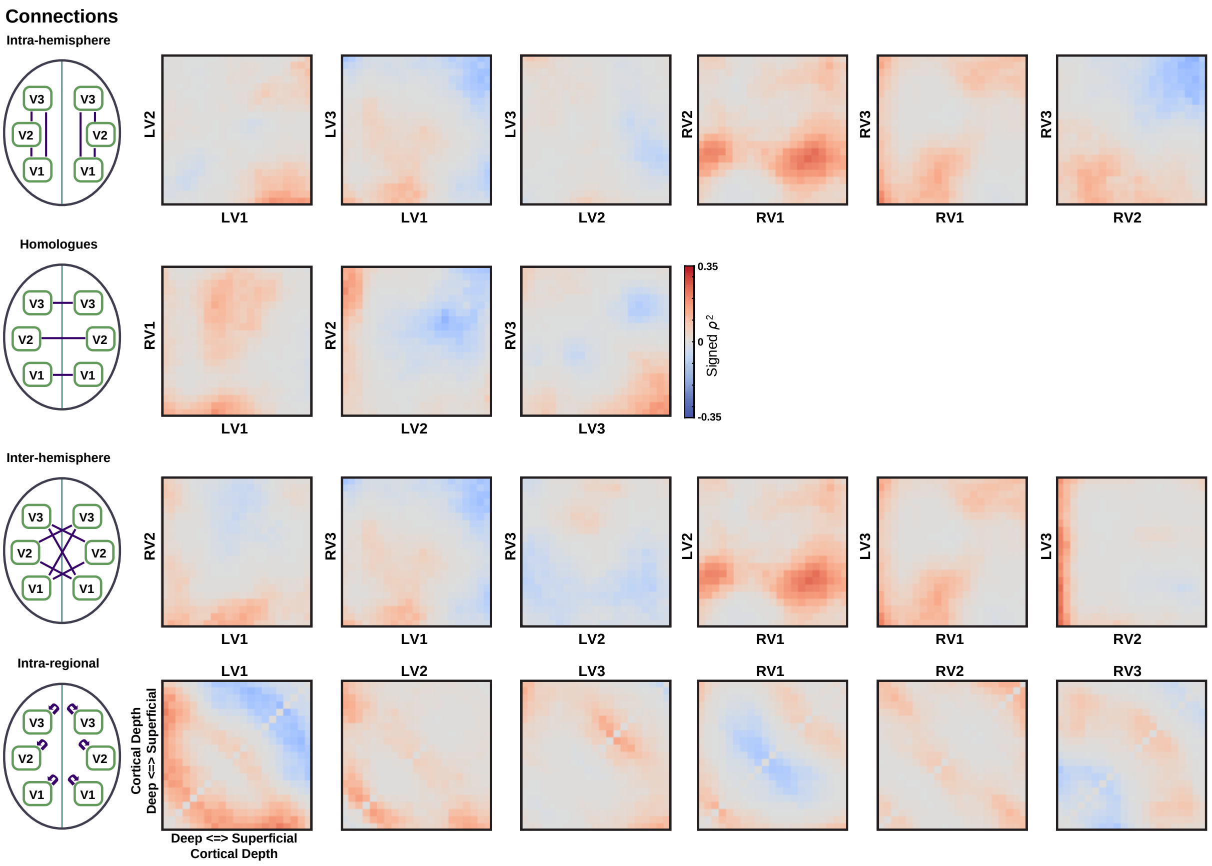


*Fig. S2: Correlation of task effects in connectivity with task effects in alpha power for individual region pairs, related to Figs. 3 and 4. Correlations are expressed as the square of the Spearman correlation multiplied by the sign of the correlation. Regions combinations are grouped as indicated in the left column.*

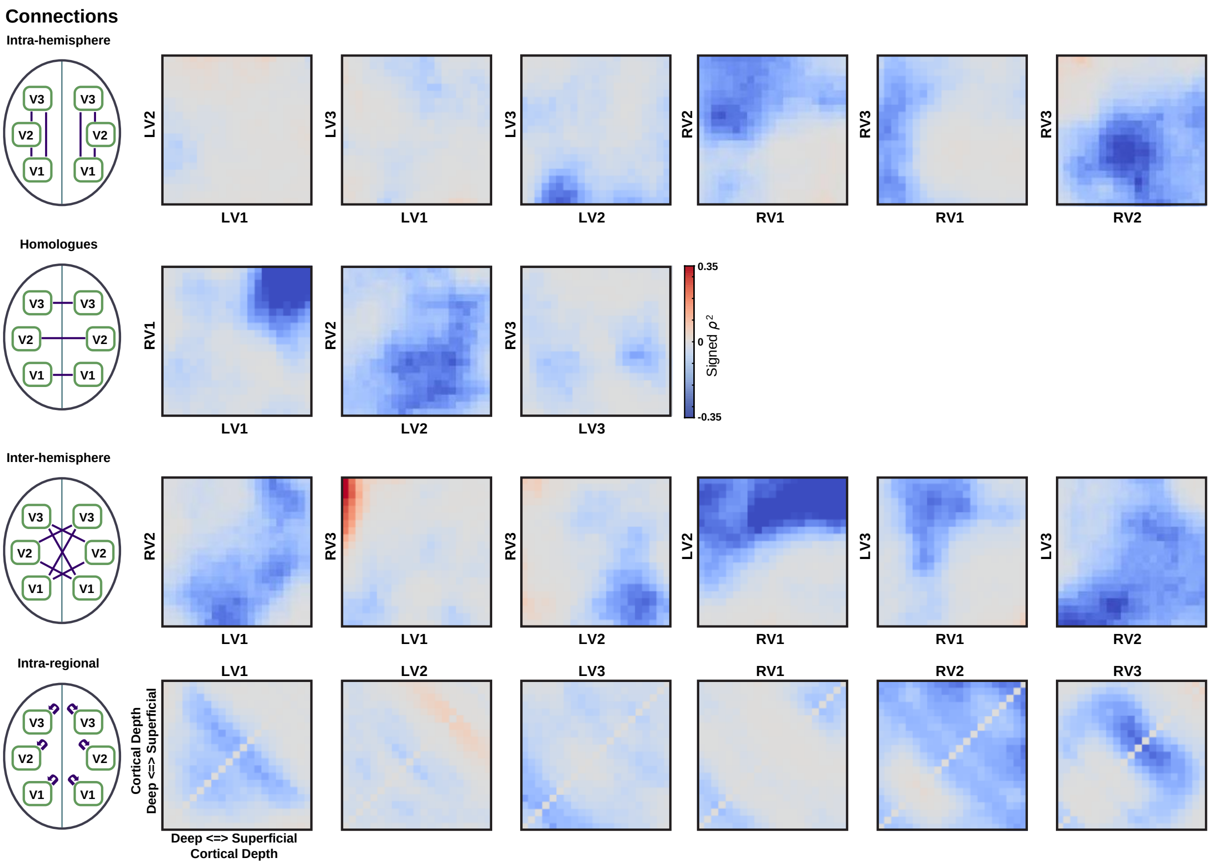


*Fig. S3: Correlation of task effects in connectivity with task effects in beta power for individual region pairs, related to Figs. 3 and 4. Correlations are expressed as the square of the Spearman correlation multiplied by the sign of the correlation. Regions combinations are grouped as indicated in the left column.*

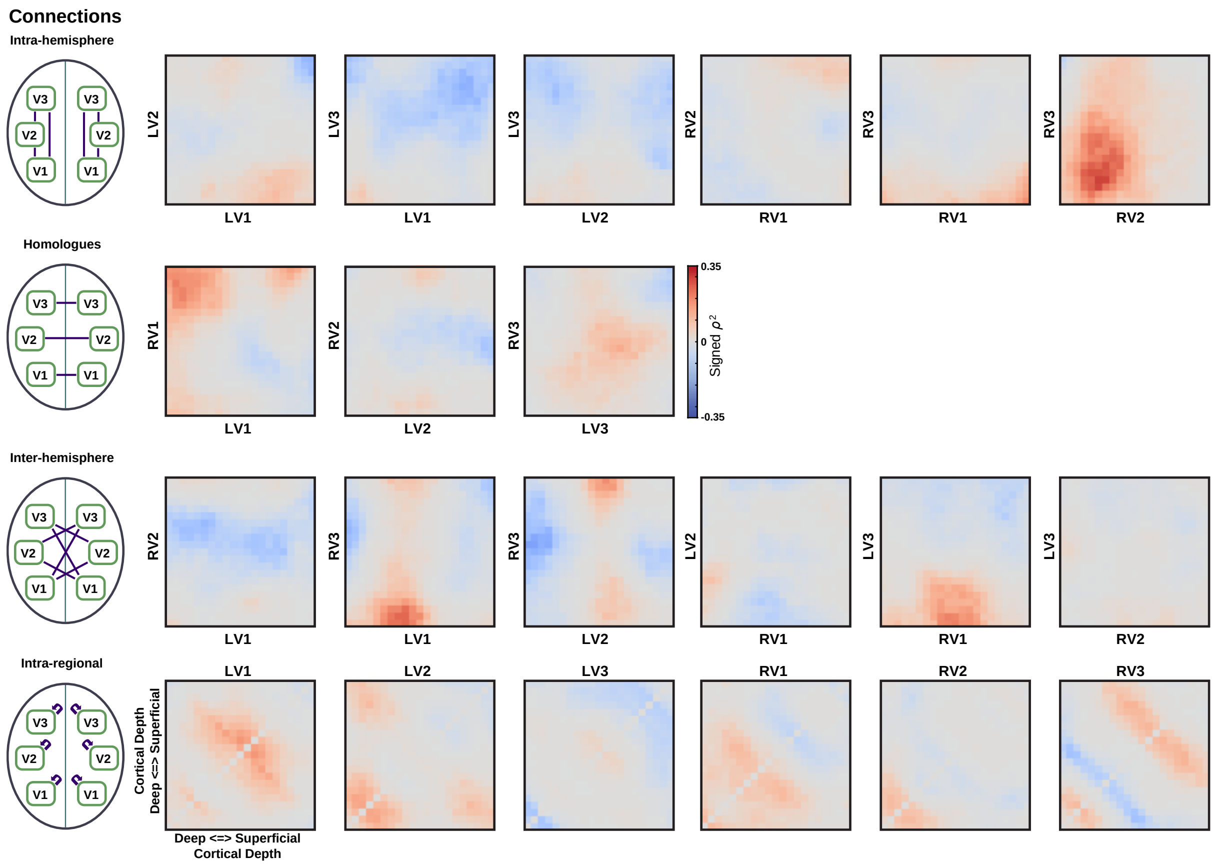


*Fig. S4: Correlation of task effects in connectivity with task effects in gamma power for individual region pairs, related to Figs. 3 and 4. Correlations are expressed as the square of the Spearman correlation multiplied by the sign of the correlation. Regions combinations are grouped as indicated in the left column.*

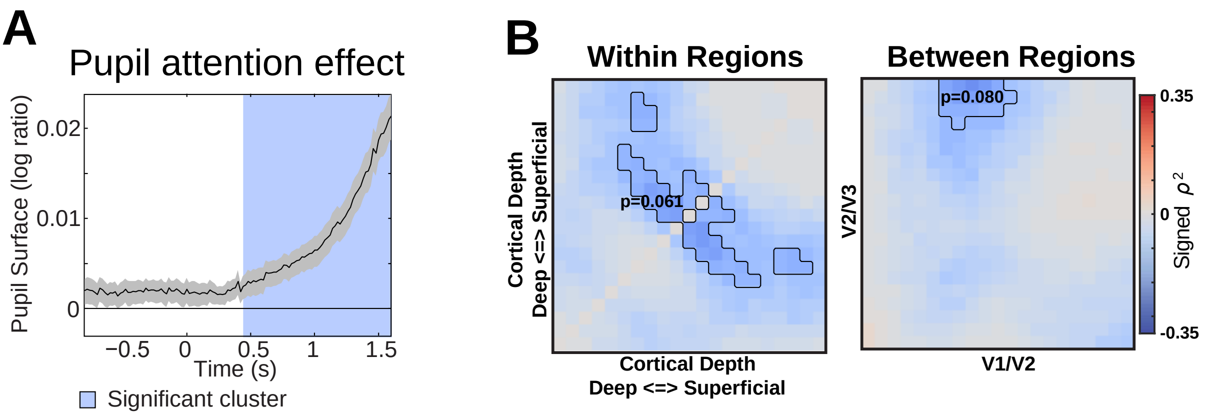


*Fig. S5: Pupil size effects. A) Attention effect for the pupil size computed as a log-ratio between the two conditions, related to Figs. 3 and 4. Adapted from Scheeringa et al. (2016). B) Signed partial Spearman correlation of pupil size with laminar connectivity, controlling for the attention effects in alpha, beta and gamma power.*

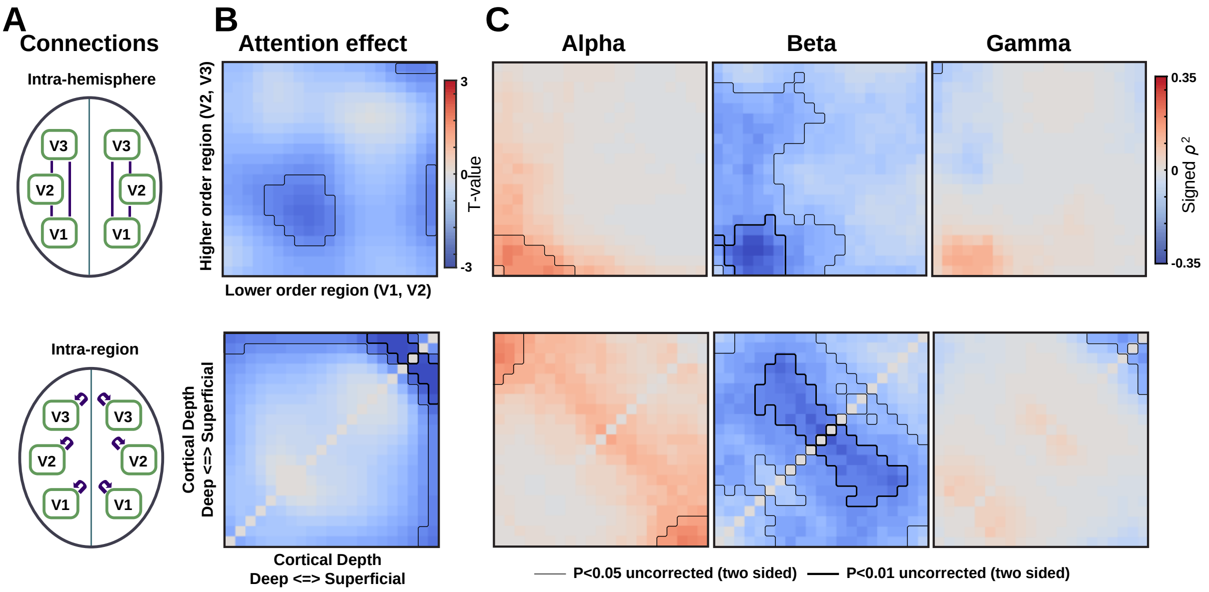


*Fig. S6: Results for the top 5% activated vertices, related to Figs. 3 and 4. A) Schematic overview of the groupings of connections between and within regions (bottom schematic) over which the effects were averaged for the results depicted at the same rows in panels B-E. B) Attention effect on laminar connectivity. C) Relation between attention effect in EEG power and laminar fMRI connectivity. Depicted is the signed squared partial Spearman correlation over subject (n=30). For each frequency band, the effects of other frequency bands and pupil size are partialed out. The p-values mentioned in panels C-D pertain to the largest supra-threshold cluster in each sub-panel and are based on a non-parametric cluster based permutation test. The results are based on the average BOLD signal in the top 5% activated vertices.*

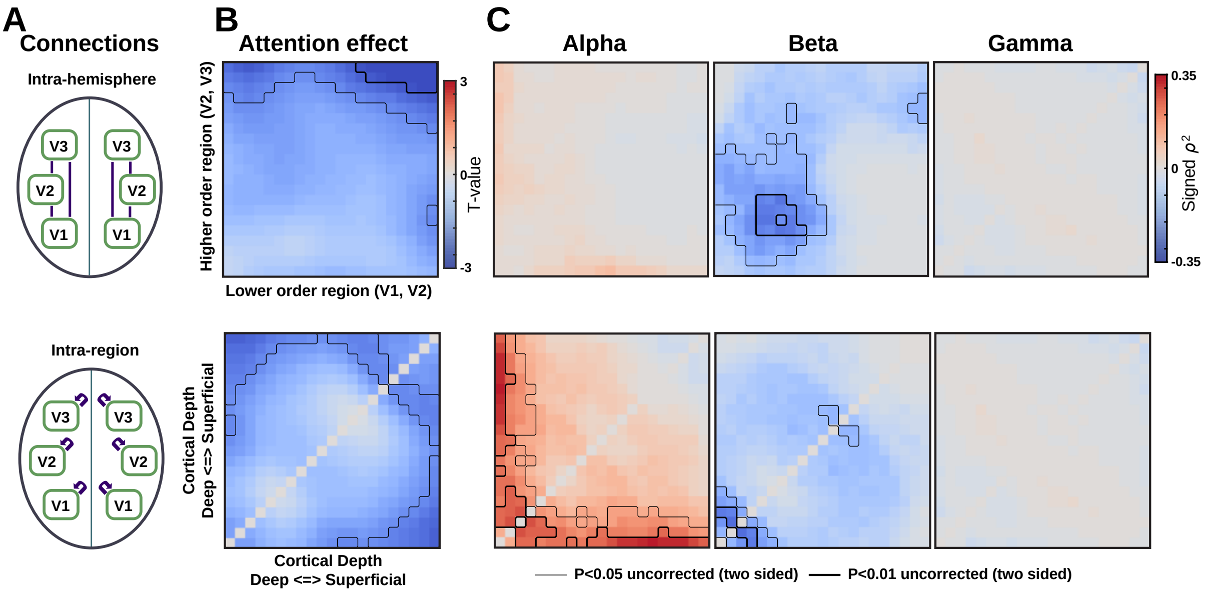


*Fig. S7: Results for the top 25% activated vertices, related to Figs. 3 and 4. Similar as for Fig. S6 but for the top 25% activated vertices.*
